# Diverse and Flexible Strategies Enable Successful Cooperation in Marmoset Dyads

**DOI:** 10.1101/2025.05.06.652115

**Authors:** Olivia C. Meisner, Weikang Shi, Amrita Nair, Gargi Nandy, Monika P. Jadi, Anirvan S. Nandy, Steve W. C. Chang

## Abstract

In humans, cooperation relies on advanced social cognition, but the extent to which these mechanisms support cooperation in nonhuman primates remains unclear. To investigate this, we examined freely moving marmoset dyads in a cooperative lever-pulling task. Marmosets successfully coordinated actions, relying on social vision rather than environmental cues. Blocking visual access or replacing the partner with an automated agent disrupted coordination. Causal dependencies between social gaze and pull actions revealed both gaze-dependent and gaze-independent strategies. Cooperation depended on social relationships, including dominance, kinship, and sex. Remarkably, marmosets adapted strategies based on partner identity, indicating rapid social learning and memory. Altogether, these findings show that flexible, cognitively driven cooperation extends more broadly across primates than previously recognized, informing our understanding of cooperative behavior’s mechanisms and evolution.

## Introduction

Cooperation is a fundamental social behavior that allows individuals to achieve shared goals, from mutual survival to group cohesion. While evidence for cooperative behavior exists across diverse taxa, from insects to apes (*1-4*), the cognitive flexibility and social sophistication driving such behavior have been primarily studied in humans and often assumed to be uniquely human (*5, 6*). Nonhuman primates are known to cooperate through predominantly kin-based mechanisms (*7*) or reciprocal exchanges (*8*), but the extent to which they develop cognitively controlled social strategies that can be adapted based on dynamic social factors remains unclear. Addressing these questions is essential for uncovering the cognitive mechanisms that underpin cooperation and understanding its evolutionary origins.

Common marmosets (*Callithrix jacchus*), a socially tolerant New World monkey species, provide a compelling model to investigate these questions. As cooperative breeders that live in family groups, they exhibit remarkable social cohesion and hallmarks of sophisticated communication, including vocal turn-taking and gaze-following (*9, 10*). While these behaviors have often been considered reflexive, recent discoveries, such as their ability to vocally label conspecifics (*11*), a capacity previously thought to be limited to humans and a few other mammalian species (*12, 13*), reveal a surprising degree of cognitive sophistication. Nonhuman primates are known to actively use gaze during strategic social interactions (*14-16*). Like other nonhuman primates, marmosets are also highly visual. However, the strategic use of social gaze in this highly cooperative species remains poorly understood. Furthermore, the ability of marmosets to develop and adapt social strategies based on relationship factors such as conspecific identity and shared experience remains unexplored.

Here we demonstrate that marmosets exhibit sophisticated cooperative abilities that extend beyond simple behavioral coordination. By combining computational approaches with naturalistic behavioral contexts, we characterize previously unquantifiable dynamics of cooperative behavior. Using an automated cooperative pulling task paired with automated gaze tracking, we reveal that marmosets rely on active social monitoring and gaze-dependent communication strategies to cooperate. Their performance is modulated by dominance, kinship, and sex, and they flexibly adapt their strategies day-to-day based on their partner’s identity and shared history. These findings expand our understanding of advanced social strategies in nonhuman primates, revealing sophisticated cooperative abilities in species evolutionarily distant from humans. This discovery provides new insights into the depth and complexity of primate social cognition, suggesting that advanced cooperative abilities may be more widespread across the primate lineage than previously recognized.

## Results

### Cooperative learning and performance improvements

We initially trained three dyads of cage-mate marmosets (two male-female pairs, one female-female pair) on a cooperative pulling task using the ‘MarmoAAP’, a recently developed marmoset apparatus for automated pulling (*17*). Each marmoset was placed in a transparent box with access to a pull lever. The marmosets could freely move around their boxes and engage with their levers spontaneously. After mastering the Self-Reward (SR) condition, where independent lever pulls earned individual rewards, marmosets progressed to cooperation training (Fig. 1A). In this phase, the dyad needed to coordinate their lever pulls within a cooperative time window (CTW) to receive rewards. Training began with a 3-second CTW (L3) and gradually decreased as they achieved 50% success rate at each phase. Once the dyad was successfully trained to coordinate pulls within a 1-second window (L1), we referred to the task as Mutual Cooperation (MC).

**Figure 1.**
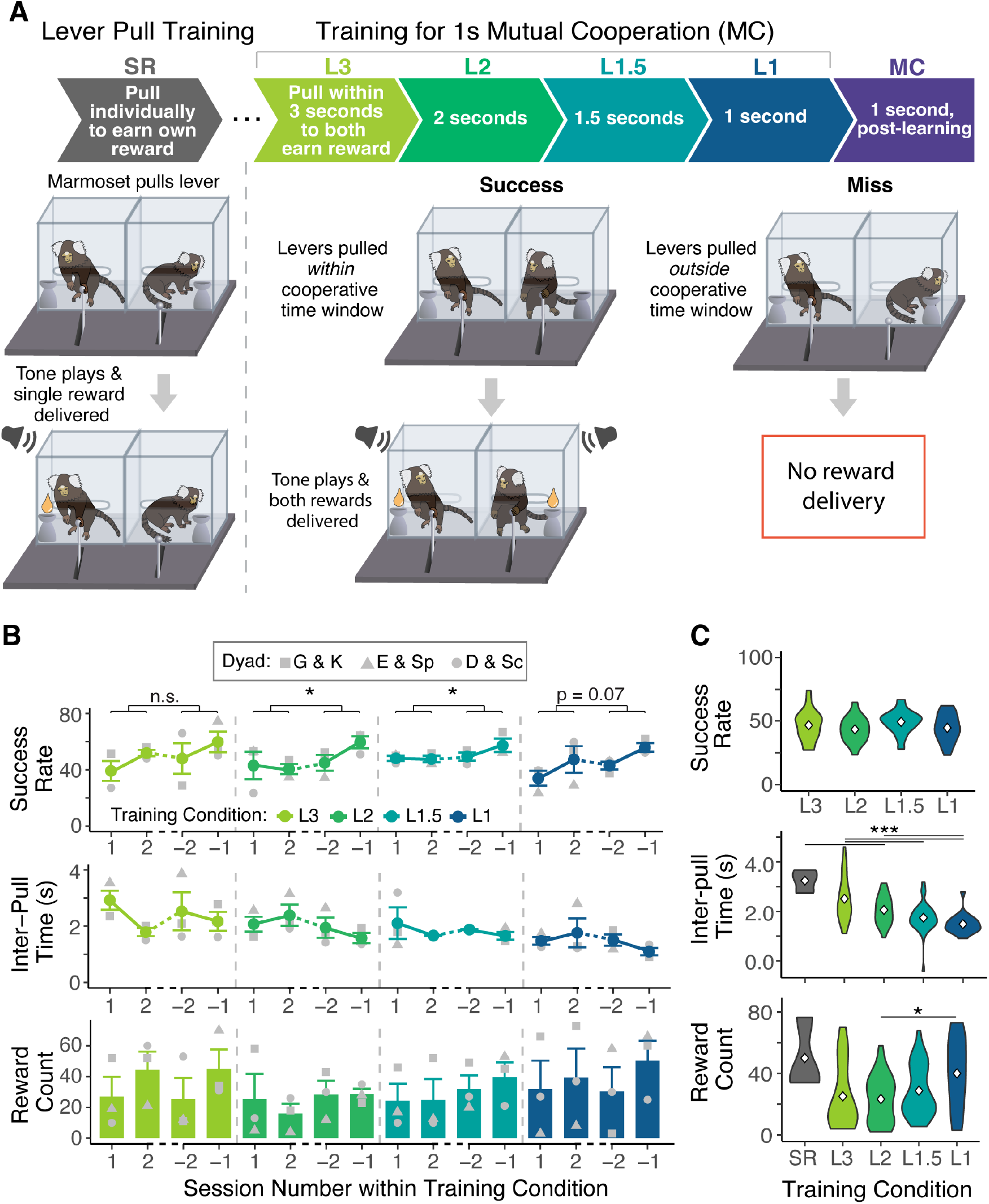
Task Structure and Learning Progression in Mutual Cooperation Training. (**A**) Schematic of Mutual Cooperation (MC) training phases. Marmosets were first trained to pull levers individually to earn rewards. They then progressed through cooperative phases with decreasing cooperative time windows (3s, 2s, 1.5s, 1s) to pull levers together for a reward. (**B**) MC learning performance metrics calculated for the first two sessions (denoted as 1, 2) and last two sessions (−1, -2) of each training phase. *Top*: Success rate calculated as number of successful lever pulls divided by total number of lever pulls of an animal in a session, averaged within a dyad (n = 2 animals), further averaged across dyads (n = 3 dyads). *Middle*: Inter-pull time calculated as time between lever pulls of a dyad in a session, averaged across all dyads (n = 3). *Bottom*: Reward count calculated as total rewards earned by a dyad per session, averaged across all dyads. (**C**) MC learning performance metrics pooled across all dyads and all sessions for each cooperation training phase. White diamond indicates the mean. Asterisks denote levels of statistical significance (*p < 0.05, ***p < 0.001). (n = 3 dyads for all plots)

We quantified three key metrics to track the dyads’ learning to cooperate across sessions: success rate (the ratio of successful lever pulls to total pulls per session), inter-pull time (the average time between lever pulls of the two marmosets within a session), and rewards earned (the total number of rewards earned by a dyad per session) (Fig. 1B-C). The marmoset dyads exhibited a reliable pattern of performance improvement from the beginning to the end of each training phase (Fig. 1B, *Top*). When they progressed to a subsequent phase with a shorter CTW, their performance typically declined at first, indicating a need to adjust to the more challenging conditions of the task. When comparing the first two sessions to the last two sessions within each training phase, dyads exhibited significantly higher success rates in the last two sessions for L2 and L1.5, with a marginally significant increase for L1 (p = 0.18 (L3), 0.02 (L2), 0.01 (L1.5), 0.07 (L1), Paired t-test; Fig. 1B *Top*). This demonstrates that marmosets adapted and improved their performance over the course of a training phase. When averaged across all sessions for each training phase from all three dyads, the marmosets showed consistent success rates across all four phases (p = 0.13, ANOVA main effect of training phases; Fig. 1C *Top*). Dyads maintained high performance levels with success rates of 45.9 ± 10.4% in L1 despite the increased difficulty of the task due to the shortening of CTWs.

Additionally, there were significant reductions in inter-pull times across the training phases (p < 0.001, ANOVA main effect of training phase; Fig. 1C *Middle*), indicating that the marmoset dyads synchronized their lever pulls more closely over time. Notably, the marmosets earned the highest number of rewards per session during the most challenging 1-second CTW (p = 0.03, ANOVA main effect of training phase; Fig. 1C *Bottom*). These findings indicate that the marmosets were effectively learning the structure and requirements of the MC task, enabling them to improve their performance with each session (Movie S1).

### Critical roles of social vision in cooperation

To further verify that the marmosets fully understood the nature of the task and were genuinely cooperating, as well as to identify the specific social interactions contributing to their successful coordination, we conducted several control experiments (Fig. 2A-C). In the No-Vision condition (NV), an opaque barrier between the marmosets prevented visual contact to both the partner and the partner’s lever (Fig. 2A), testing whether successful cooperation could be achieved through random frequent pulling or through use of non-visual cues like vocalizations or rhythmic pulling. Compared to MC, the NV condition showed significantly decreased success rates (p < 0.001, ANOVA main effect of task type; MC vs. NV p < 0.001, post hoc Tukey HSD), increased inter-pull times (p < 0.001, ANOVA main effect of task type; MC vs. NV p < 0.001, post hoc Tukey HSD), and reduced rewards earned (p < 0.001, ANOVA main effect of task type; MC vs. NV p < 0.001, post hoc Tukey HSD), demonstrating that the success in the MC condition was not merely due to frequent lever pulling (Fig. 2D-E). Cross-correlation analysis of dyad pull times revealed impaired coordination in NV, with lower peak cross-correlation coefficients (p < 0.001, ANOVA main effect of task type; p < 0.001, post hoc Tukey HSD MC vs. NV; Fig. 2F-G) and more variable peak lags (p < 0.001, ANOVA main effect of task type; p < 0.001, post hoc Tukey HSD MC vs. NV; Fig. 2F-G). The observed cross-correlation coefficients exceeded that of the shuffled data, indicating that the simultaneous pulling in MC was not due to chance. These results demonstrate that marmosets relied on visual information for successful cooperation (Movie S2).

**Figure 2.**
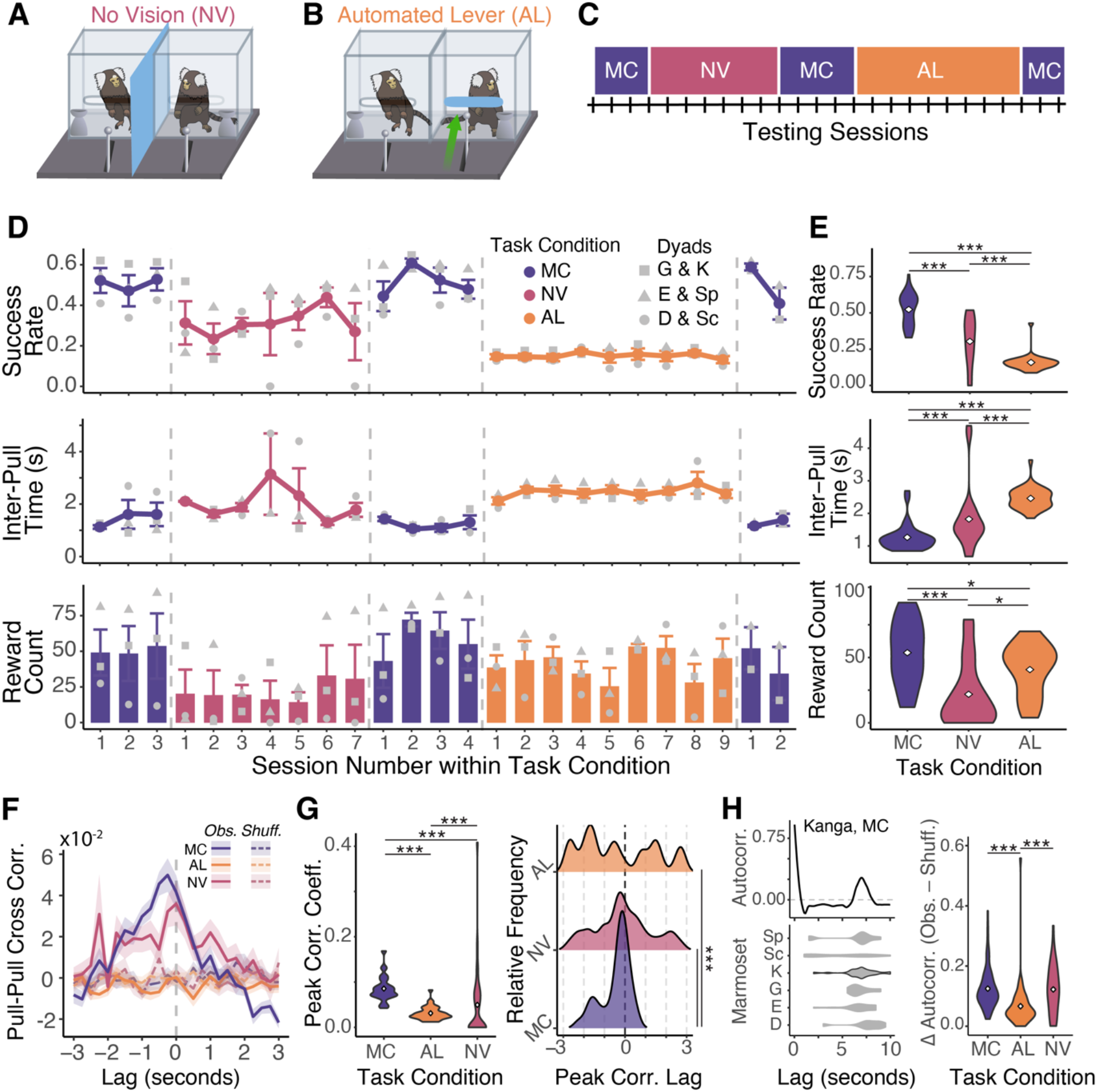
Role of social vision in cooperative interactions. (**A**-**B**) Illustrations of the No-Vision (NV) (**A**) and Automated-Lever (AL) (**B**) control conditions. (**C**) Example timeline of interleaving the 1s Mutual Cooperation (MC) condition with the NV and AL control conditions. (**D**) Session-wise performance metrics for the MC, NV, and AL conditions. *Top*: Success rate calculated as number of successful lever pulls divided by total number of lever pulls of an animal in a session, averaged within a dyad (n = 2 animals), further averaged across dyads (n = 3 dyads). *Middle*: Inter-pull time calculated as time between lever pulls of a dyad in a session, averaged across all dyads (n = 3). *Bottom*: Reward count calculated as total rewards earned by a dyad per session, averaged across all dyads. (**E**) Performance metrics pooled across all dyads and all sessions within each task condition. White diamonds indicate the means. (**F**) Cross-correlation coefficient values between pull times, across task conditions. They were computed per dyad per session, using dominant animal as the reference, and averaged across dyads and sessions for both observed and shuffled data. (**G**) Peak correlation coefficient across task conditions. *Left*: Violin plots of peak correlation values from each session by task condition. *Right*: Distribution of lag values at peak correlation from each session by task condition. White diamonds indicate the means. (**H**) Temporal structure of lever-pulling behavior. *Top left*: Example autocorrelation function from one marmoset (Kanga) during an MC session. *Bottom left*: Distributions of the lag times at peak autocorrelation for each marmoset (Sp, Sc, K, G, E, D) during MC sessions. *Right*: Difference between observed and shuffled autocorrelation peaks for MC, AL, NV. Asterisks denote levels of statistical significance (*p < 0.05, ***p < 0.001). (n = 3 dyads for all plots)

To test if marmosets relied on general visual cues rather than specifically social ones, we conducted the Automated-Lever control experiment (AL; Fig. 2B). In this setup, one marmoset was blocked from accessing their lever and instead had their lever operated by a computer program to mimic typical marmoset pulling frequency in full view, while the other retained control of their lever and could earn rewards for both marmosets by pulling within the CTW relative to the automated lever’s movement. Compared to MC, the AL condition led to significantly decreased success rates (p < 0.005, post hoc Tukey HSD), increased inter-pull times (p < 0.001, post hoc Tukey HSD), and fewer rewards earned (p = 0.03, post hoc Tukey HSD) (Fig. 2D-E). Cross-correlation analysis of dyad pull times revealed the lowest coordination in AL, with reduced peak correlation coefficients (p < 0.001, post hoc Tukey HSD) and highly variable peak lags (p < 0.001, post hoc Tukey HSD) (Fig. 2F-G). Even after ten consecutive sessions (Fig. 2D orange), marmosets did not develop an alternative strategy based on viewing automated lever movements, emphasizing that they specifically relied on social visual cues rather than general visual information, and that reactively timing the pulls to the automated lever movements was challenging (Movie S3).

Given that the dyads were able to maintain moderate success rates on the NV task, we hypothesized they could employ a gaze-independent strategy involving rhythmic coordination to facilitate successful lever pulls. To test this *‘pull-in-rhythm*’ strategy, we analyzed autocorrelation patterns in individual marmosets’ pulling behavior (Fig. 2H). This analysis revealed consistent rhythmic pulling across marmosets with peaks typically occurring at 5-10 seconds lag (Fig. 2H, *Bottom Left*). Comparing the strength of these temporal patterns across task conditions, we found significantly stronger rhythmic structure in both MC and NV conditions compared to AL (p < 0.001, ANOVA main effect of task type; p < 0.001, post hoc Tukey HSD; Fig. 2H, *Right*). This provides direct evidence that marmosets employ a strategy of maintaining a specific rhythmic pulling pattern to facilitate their cooperation.

Lastly, given that marmosets are well known to use vocalizations to communicate with conspecifics (*18*), we quantified the occurrences of different call types when marmosets were participating in the cooperative and control conditions (Fig. S1). While there were no differences in chirp calls across conditions (Fig. S1A), NV had significantly more phee calls than other conditions (Fig. S1B). We further analyzed the timing of these phee calls during NV, distinguishing between active periods (when marmosets were pulling levers) and rest periods (when they were not pulling) (Fig. S1C). Most phee calls occurred during rest periods rather than during active task performance (p = 0.038, paired t-test, n = 21 sessions) suggesting they were not being used to coordinate lever pulls. This aligns with the function of phee calls as long-distance contact calls, likely produced more frequently in NV because marmosets were experiencing social isolation when unable to see each other.

### The use of social gaze in cooperative interactions

The performance deficits in the NV and AL conditions provided evidence that the marmosets relied on social vision to cooperate. We further hypothesized that marmosets strategically used social gaze to the partner to collect task-relevant information and facilitate cooperation. To test this, we tracked six facial features of the freely moving marmosets to identify their social gaze (Fig.3A; Movie S4), which is predominantly accomplished by head movements in marmoset natural behaviors (*19*), and quantified the rate of social gaze (total number of gazes divided by the session time). We found an adaptive use of social gaze based on multiple pieces of converging evidence. First, over the course of training from the SR to MC, we observed an increased rate of social gaze (MC vs. SR p < 0.001, L1 vs. SR p < 0.01, L1 vs. L2 p < 0.01, post hoc Tukey HSD; Fig. 3B). The social gaze rate also dropped in the NV condition (Fig. S2). These results suggest that marmosets increase their use of social gaze, specifically in the MC condition. Second, when we additionally quantified the times when marmosets looked at the partner’s lever (partner lever gaze), the marmosets gazed more frequently at the partner’s levers than at their partners, specifically when the partners were assigned the automated lever (partner AL vs. SR p = 0.011, partner AL vs. MC p < 0.01, partner AL vs. self AL p < 0.001, post hoc Tukey HSD; Fig. 3C). This suggested that marmosets adapted their gaze strategy to the automated lever movement. However, low success rates during AL (Fig. 2D,E) indicate that reading the partners’ actions, rather than reacting to the lever movement alone, was essential for successful cooperation.

**Figure 3.**
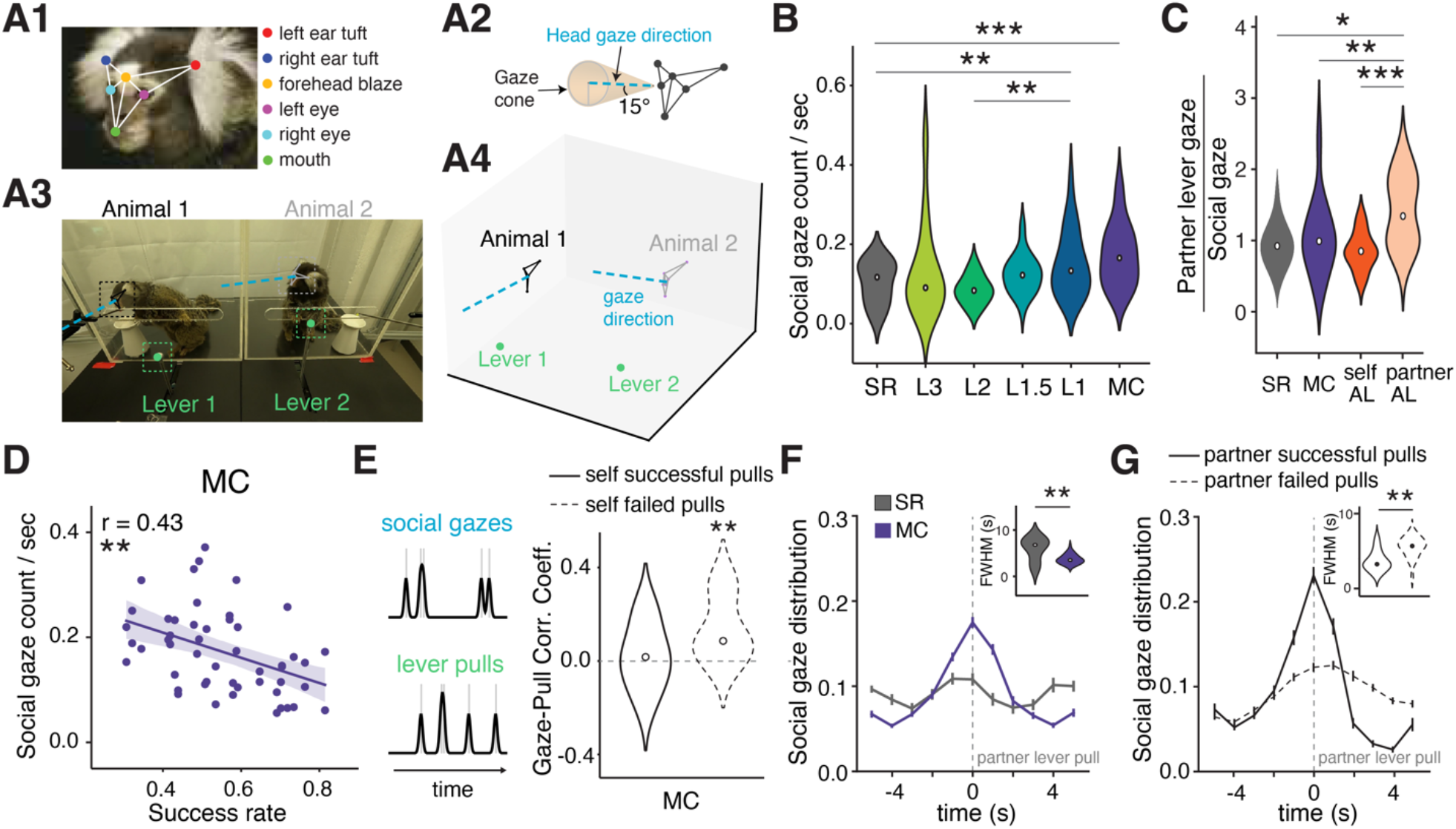
Social gaze dynamics during cooperative interactions. (**A**) Analysis pipeline for tracking freely moving marmosets. We tracked six facial parts using DeepLabCut (*20*) (**A1**). Gaze direction was defined as a 15-degree gaze cone perpendicular to the facial plane (**A2**). We triangulated facial parts from multiple camera viewpoints using Anipose (*21*) for 3D tracking (**A3, A4**). Social gaze was defined as the event when one animal’s gaze cone intersected with another animal’s face frame. (**B**) Social gaze rate in each training condition (n = 6 animals for SR to L1, n = 10 animals for MC). (**C**) Partner-lever gaze to social gaze ratio in SR, MC and AL control conditions (n = 8 animals). (**D**) Relationship between social gaze rate and success rate during MC (n = 25 sessions x 2 animals). (**E**) Correlation between social gaze epochs and self-lever pull epochs during MC (n = 10 animals). (**F**) Social gaze distribution aligned to the partner’s lever pull time in SR and MC (n = 10 animals). Inset shows the full width at half maximum (FWHM) of each distribution. (**G**) Social gaze distribution in MC aligned to the partner’s successful and failed lever pull times (n = 10 animals). Asterisks denote levels of statistical significance (*p < 0.05, **p < 0.01, ***p < 0.001): ANOVA and post-hoc Tukey’s HSD test for panel B and C; linear regression for panel D; Wilcoxon test and Mann-Whitney U test for panel F and G.

To understand the functionality of social gaze in cooperative behavior, we focused on the MC condition, where social gaze was most prevalent. Social gaze counts were negatively correlated with success rate (Fig. 3D), suggesting reduced reliance on social gaze during sessions with higher cooperative performance. We thus hypothesized that marmoset dyads use more social gaze during failed cooperative pulls to exchange information that facilitates the resynchronization of lever pulls to restore the ‘*pull-in-rhythm*’ strategy of cooperation. We tested this hypothesis by correlating temporal periods of gazes and pulls, session by session. We found a positive correlation between social gaze and failed pulls epochs (p < 0.01, Wilcoxon signed-rank; Fig. 3E), but not between social gaze and successful pulls (p = 0.6, Wilcoxon signed-rank; Fig. 3E). These temporal relations suggest that marmoset dyads are likely to use social monitoring as a strategy to re-establish rhythmic pulling after unsuccessful coordination attempts.

Next, we examined the precise timing of social gaze in relation to lever-pulling behaviors. In MC, social gaze distribution peaked around the partner’s pulls (Fig. 3F) with a significantly stronger temporal alignment in MC than in SR (full width at half maximum (FWHM) of distribution in MC vs. SR, p < 0.01, Mann-Whitney U test; Fig. 3F *inset*), indicating that social gaze was temporally coupled to the partner’s action in the MC. Interestingly, social gaze was more temporally anchored to the partner’s successful lever pulls than the partner’s failed pulls (FWHM of distribution in successful vs. failed pulls, p < 0.01, Mann-Whitney U test; Fig.3G *inset*). Together, the pairwise pull-gaze analyses revealed two distinct modes of social monitoring: a precise, temporally coupled mode during successful coordination and a broader, more vigilant mode when coordination breaks down. These results suggest the presence of a ‘*gaze-and-pull*’ strategy of cooperation in addition to the previously identified ‘*pull-in-rhythm*’ strategy.

To further investigate cooperation strategies, we systematically analyzed the causal relationships within the multivariate marmoset dyad behavior data and fit predictive models to test our strategy hypotheses.

### Diverse social strategies used in cooperative interactions

To assess the composition of marmoset cooperative strategies, we used generative Dynamic Bayesian Networks (DBN), a class of probabilistic graphical models describing causal dependencies, to predict the behavioral dynamics of marmosets during MC (see Methods). We trained two types of models using the multivariate behavior data: a full model incorporating learned causal dependency graph (Fig. 4A1) and models with causal dependency graphs describing the hypothesized strategies suggested by our previous analyses (Fig. 4A2). The ‘*pull-in-rhythm*’ strategy would be reflected as enhanced dependencies between the two animals’ pulls (pull-to-pull dependency) (Fig. 4A2). The ‘*gaze-and-pull*’ strategy would be reflected in two distinct types of dependency dynamics: social-attention dependency (partner’s lever pull triggers a marmoset’s own social gaze) and gaze-lead-pull dependency (marmoset’s own social gaze leads to its own lever pull).

**Figure 4.**
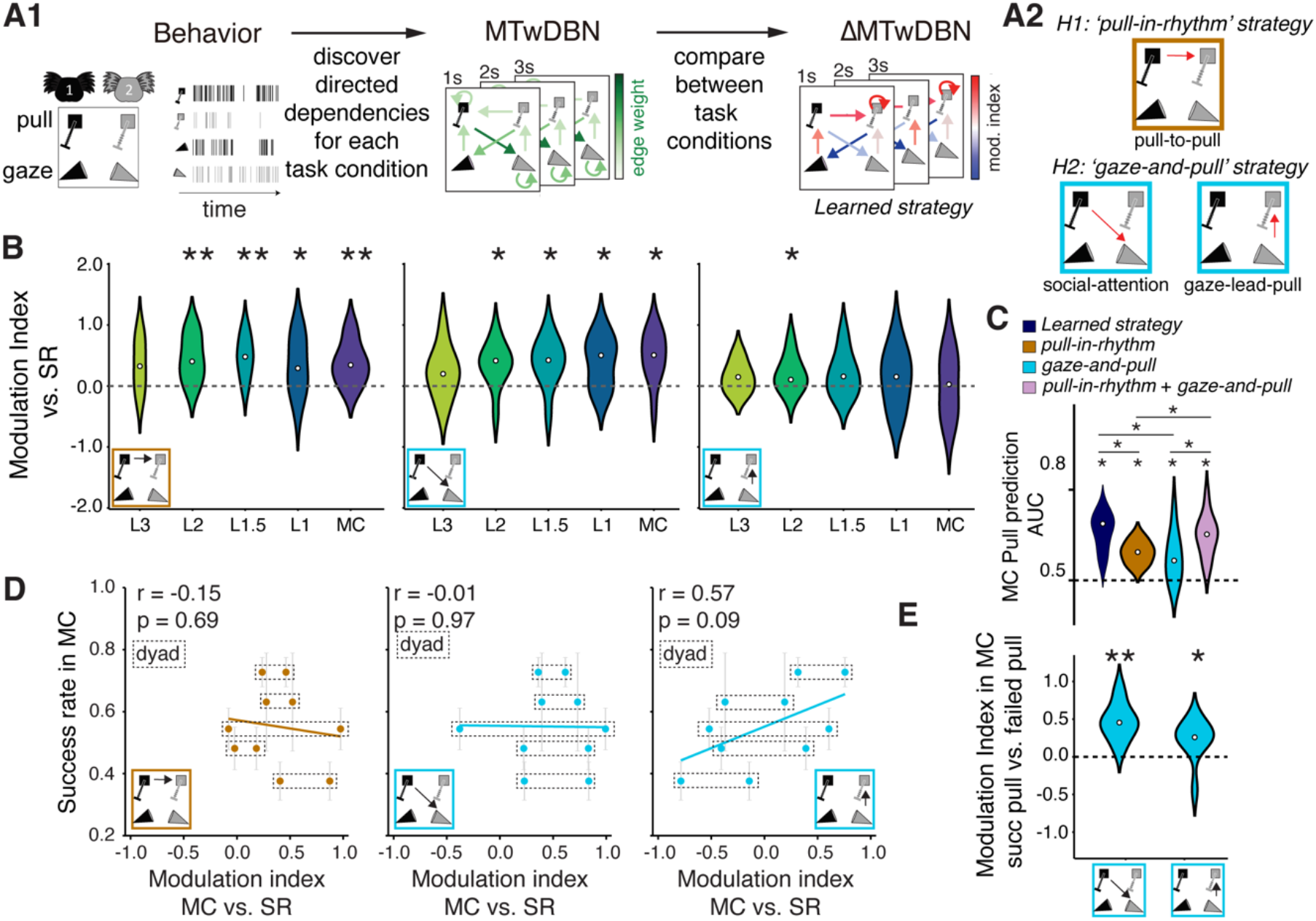
Social strategies in the cooperative pulling task. (**A1**) Analysis pipeline for discovering social strategy. A multi-timelag weighted Dynamic Bayesian Network (MTwDBN) was used to discover directed dependencies among four behavioral events: social gaze and lever pull of two animals. We discovered the cooperation strategy of the dyad by comparing the dependencies between cooperation conditions [learning phases L3, L2, L1.5, L1, and Mutual Cooperation (MC)] and Self Reward (SR) and calculating their modulation indices (∆MTwDBN). (**A2**) The two hypothesized strategies expressed as DBN graphs: ‘*pull-in-rhythm’* strategy involving pull-to-pull dependencies, and ‘*gaze-and-pull’* strategy involving both the gaze-lead-pull dependencies and social-attention dependencies. (**B**) Modulation indices for cooperation conditions from L3 to MC (n = 6 animals for L3 to L1, n = 10 animals for MC), when compared with SR. (**C**) MC pull prediction performance. DBN predictive models with different dependency structures were used for prediction. The “learned structure” model contained all significant dependencies from the ∆MTwDBN fit to MC and SR data (A1); ‘*pull-in-rhythm’* and ‘*gaze-and-pull’* strategy models had dependencies as described in (A2). Dashed line indicates performance at chance. (**D**) Dependency modulation vs success rates. (**E**) Dependency modulation across partner’s successful vs failed pulls during MC. Asterisks denote levels of statistical significance (*p < 0.05, **p < 0.01): Wilcoxon signed-rank test for panel B, C and E; linear regression for panel D. (n = 10 animals from 5 dyads for panels C to E).

To estimate causal dependency graphs among the social gazes and lever pulls of the dyad at multiple time lags and under all task conditions, we fit multi-timelag weighted Dynamic Bayesian Network (MTwDBN) models to behavior data (Fig. 4A1, Fig. S3A). To isolate strategies that were unique to cooperation, we compared the DBN graphs of cooperative conditions (training phase L3 to L1 and MC) with SR and estimated the modulation graph ∆MTwDBN (Fig. S3B, see Methods). We analyzed the ∆MTwDBN graphs for these three dependencies (pull-to-pull, social-attention, and gaze-lead-pull) to evaluate their modulation during cooperation (Fig. S4 and Movie S5).

During cooperation, we found comparable dependency modulations across all the time lags in our model (Fig. S5), and hence considered lag-averaged modulation indices for further analysis (Fig. 4B). Both pull-to-pull dependencies and social-attention dependencies were enhanced compared to SR across later training phases, indicating a consistent use of both the ‘*pull-in-rhythm’* and ‘*gaze-and-pull’* strategy once the marmosets learned the task contingency in L2. Importantly, these behavioral patterns were not present at the beginning of training (L3) but emerged as marmosets learned the task, suggesting that these strategies developed specifically to meet the demands of cooperative coordination rather than representing pre-existing behavioral tendencies. While the partner monitoring behavior, as measured by the social-attention dependency, appeared to be a persistent component of the ‘*gaze-and-pull’* strategy, the gaze-lead-pull dependency was also enhanced in L2, suggesting that marmosets also use social gaze to guide their cooperative pulls, but only significantly so when a long cooperation window allows them to do so, i.e. L2 (Fig. 4B).

To assess the functional significance and the strategic composition of the dependencies modulated during cooperation, we used generative DBN models to predict the behavioral dynamics of marmosets during MC (see Methods). DBN models were trained with four types of dependency structures: the MC vs. SR dependency modulation graph learned from the data (full model, *learned strategy*), the hypothesized ‘*pull-in-rhythm*’ strategy graph, the hypothesized ‘*gaze-and-pull’* strategy graph, and lastly, a graph combining both strategy hypotheses (Fig. 4A2, Fig. S4). We then quantified the prediction performance of these four models on held-out data for pull actions during MC. The full model (*learned strategy*) predicted marmoset pull actions significantly better than chance (AUC = 0.64 ± 0.06) and provided an upper bound for the predictive performance. The individual strategy models all predicted marmoset pull actions significantly better than chance (Fig. 4C, AUC = 0.64 ± 0.06, 0.57 ± 0.04, 0.56 ± 0.07, 0.62 ± 0.06, respectively), with the full model outperforming the individual models (‘*learned strategy*’ vs ‘*pull-in-rhythm*’ p = 0.02, ‘*learned strategy*’ vs ‘*gaze-and-pull*’ p = 0.03, post hoc pair-wise Wilcoxon with multiple comparison correction). However, the prediction performance of the combined strategy model was comparable to that of the full model (p = 0.6, post hoc pair-wise Wilcoxon with multiple comparison correction). These results indicate that the dependency modulations and strategies identified using our approach capture the underlying structures of the social behavioral dynamics during cooperative interactions.

Having tested our hypotheses, we next examined how dependency strength influences cooperation success. The modulation indices for pull-to-pull dependencies and social-attention dependencies showed no correlation with success rate. However, the modulation index of the gaze-lead-pull dependencies was weakly positively correlated with success rate (p = 0.09, Spearman correlation; Fig. 4D). Moreover we found significant enhancements of both social-attention (modulation index = 0.50 ± 0.18) and gaze-lead-pull (modulation index = 0.20 ± 0.29) dependencies during successful compared to failed pulls, indicating greater deployment of the *‘gaze-and-pull’* strategy (Fig. 4E). Altogether, these results demonstrate that the marmoset dyads developed and utilized both gaze-independent and gaze-dependent strategies to facilitate cooperation, with task performance influenced by the strength of these strategies, particularly the gaze-dependent ones.

### Impact of social relationships on cooperative interactions

To understand how social relationships influence cooperation, we examined how three social factors –dominance, kinship, and sex – shaped cooperative behaviors. Using established dominance criteria to define hierarchy in our dyads (Methods), we examined how dominance influenced the development of cooperative pulling behaviors through cross-correlation analysis of lever pulls (Fig. 5A). With dominant marmosets’ pulls serving as the reference, negative lags indicated that the dominant’s pulls follow the subordinate’s pulls, while positive lags indicated the opposite direction.

**Figure 5.**
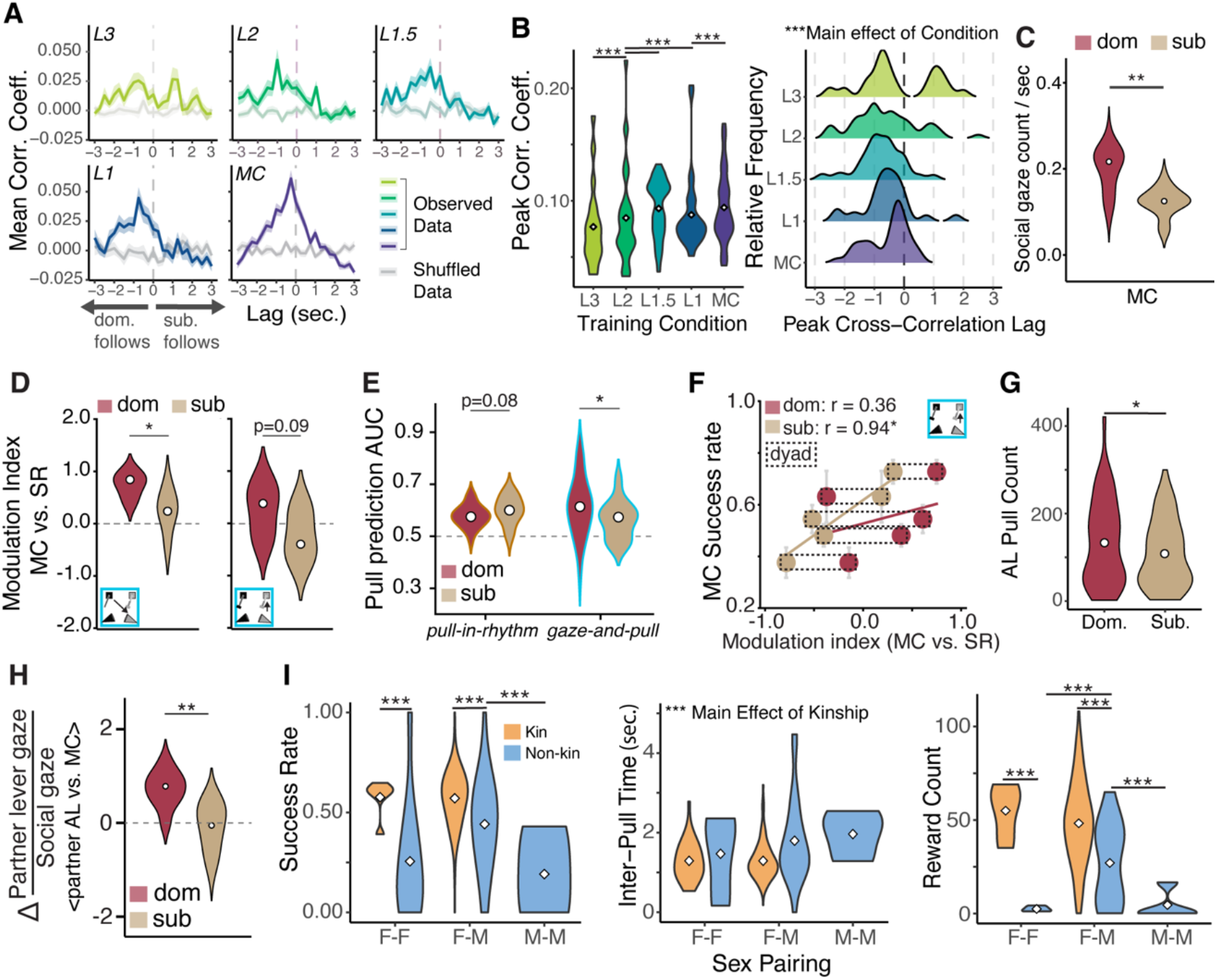
Influence of social factors on cooperative strategies. (**A**) Cross-correlation coefficient values by lag for observed and shuffled data by training phase. (**B**) Peak correlation (*Left*): Violin plots of peak correlation values from each session by training phase. Peak Lags (*Right*): Distribution of lag values at peak correlation for each session by training phase. (**C**-**H**) Influence of social hierarchy (dominant vs. subordinate) on social gaze rate in MC (**C**); modulation indices of MC for the dependencies from the ‘*gaze-and-pull*’ strategy (**D**); MC pull prediction performance of the ‘*pull-in-rhythm’* and ‘*gaze-and-pull’* strategy models (**E**); correlation between modulation index and success rate for the gaze-lead-pull dependency in MC versus SR conditions (**F**); performance on automated lever (AL) condition (**G**); and partner-lever gaze to social gaze ratio difference between partner AL and MC conditions (**H**). (**I**) Performance of mixed sex and kinship dyads (non-kin dyad counts: 4 F-F, 5 M-F, 3 M-M; kin dyad counts: 1 F-F, 3 M-F). Additionally, we included data from one more kin pair that learned the MC condition after the initial learning experiment for a total of 4 kin pairs (1 female-female, 3 male-female). Performance metrics (*Left*: success rate, *Middle*: inter-pull time, *Right*: reward count) were calculated by session for kin and non-kin female-female, female-male, and male-male (non-kin only) dyads. Metrics were calculated as described in Fig. 2D. Asterisks denote levels of statistical significance (*p < 0.05, **p < 0.01, ***p < 0.001).

Coordination between animals became progressively stronger and more temporally aligned as training advanced. The strongest coordination emerged in the final training phases compared to earlier phases (p < 0.001, ANOVA main effect of training condition; all comparisons except L1 vs. L1.5 significant at level of p < 0.001, post hoc Tukey HSD; Fig. 5B, *Left*). More importantly, we discovered a consistent shift in the timing relationship between partners’ actions. While dyads initially showed variable timing patterns of pulls, they developed a stable dynamic where dominant individuals reliably followed the actions of subordinate animals (Fig. 5B, *Right*). This emergence of a highly consistent leader-follower pattern, with dominants consistently following subordinates’ lever pulls, was remarkably consistent across all three dyads (Fig. S6), revealing that dominant marmosets adapted their pull timing to the subordinate partners’ initiations during cooperative problem-solving.

Having established that dominant marmosets adapted to follow their subordinate partners during cooperation, we examined how dominance shaped other aspects of cooperative interactions among kin pairs. Dominant animals showed higher rates of social gaze in the MC condition (p < 0.01, Mann-Whitney U test; Fig. 5C), and analysis of gaze-dependent strategies revealed that dominant animals employed the ‘*gaze-and-pull*’ strategy at a significantly higher level than subordinates (social-attention dependency p = 0.02, gaze-lead-pull dependency p = 0.09, Mann-Whitney U test; Fig. 5D). Generative DBN models more accurately predicted the pull actions of dominant animals using the ‘*gaze-and-pull’* strategy graph (p = 0.03, Mann-Whitney U test; Fig. 5E, *Right*). Conversely, the models showed slightly better prediction performance for subordinate animals when using the ‘*pull-in-rhythm*’ strategy graph (p = 0.08, Mann-Whitney U test; Fig. 5E, *Left*), suggesting distinct strategy preferences between dominant and subordinate animals. Additionally, only in subordinate animals did the strength of gaze-lead-pull dependency positively correlate with success rate (sub p = 0.04, Spearman correlation; Fig. 5F). This suggests that while dominant animals were more actively engaged overall, perhaps to closely follow subordinates’ behaviors, successful cooperation critically depended on subordinates’ strategic behaviors. Dyads achieved higher success rates when subordinate animals made greater use of the ‘*gaze-and-pull’* strategy, likely because dominant animals were consistently engaged regardless of task demands. On the other hand, subordinates’ limited use of the ‘*gaze-and-pull’* strategy became a limiting factor or ‘bottleneck’ for successful cooperation. In the AL control condition, dominant animals performed more lever pulls (p = 0.03, Mann-Whitney U test; Fig. 5G), indicating greater task engagement. They also showed stronger gaze adaptation in the AL control condition, with more frequent gaze shifts toward their partner’s automated lever, supporting a more pronounced adjustment of gaze strategy in dominant individuals (p < 0.01, Mann-Whitney U test; Fig. 5H).

To investigate how social relationship factors beyond dominance influence cooperation in marmoset dyads, we examined performance across different dyad types by creating new pairs varying in kinship and sex composition. Notably, while non-kin cooperation is traditionally considered rare in nonhuman animals (*23*), non-kin marmoset pairs demonstrated substantial cooperative ability even with only three days of interaction, though kin pairs still showed better performance across all measures (higher success rates, shorter inter-pull times, and greater reward counts; all p < 0.001, ANOVA, main effect of kinship; Fig. 5I). Among non-kin pairs, we found a significant interaction between kinship and sex pairing (success rate p = 0.04; reward count p = 0.03, ANOVA), with male-female dyads earning more rewards than both female-female (p < 0.01, post hoc Tukey HSD) and male-male pairs (p = 0.04, post hoc Tukey HSD), and achieving higher success rates than male-male pairs (p = 0.013, post hoc Tukey HSD). With respect to vocalizations, non-kin same-sex dyads showed distinct stress responses: female-female pairs exhibited increased negative vocalizations suggesting territorial and aggressive reactions (e.g., chatter, tsik, ek, tsik-ek) (*18*), while male-male pairs showed increased phee calls indicating social anxiety from separation (*24*) (Fig. S7). These findings demonstrate the profound impact of social dynamics on cooperative success, where preexisting social bonds and mixed-sex partnerships facilitate cooperation, while same-sex competition and social stress hinder it. Moreover, the substantial cooperation achieved by non-kin pairs, especially the female-male pairs, suggests that the natural ecology of a species, rather than inherent cognitive or motivational limitations, may determine whether non-kin cooperative behaviors emerge in natural settings.

### Flexible implementation of social strategies based on partner identity

Cooperative strategies in socially sophisticated species should be adaptable and tailored to the specific identity and behavior of a partner. To directly test whether marmosets develop and flexibly deploy partner-specific cooperation strategies, we had two test marmosets (Kanga and Ginger) alternate between different partners over 10 consecutive days of Mutual Cooperation (MC) (Fig. 6A). We tested flexible cooperative strategies in three dyads: Kanga/Ginger (female-female mother-daughter), Ginger/Dodson (female-male pair-mate), and Kanga/Dannon (female-male pair-mate). Strikingly, marmosets exhibited clear, partner-specific behavioral patterns that varied systematically with the identity of their partner on a given session (Fig. 6B-C and Movie S6). For instance, Ginger showed systematic variations in multiple behavioral measures, including pull ratio, inter-pull interval, social gaze frequency, and social-attention dependency, that depended on her partner’s identity (Fig. 6B). Additionally, her pulling behavior was distributed differently in relation to her partner’s lever pulls, showing distinct probability patterns depending on the partner (p < 0.001, Mann-Whitney U test; Fig. 6C).

**Figure 6.**
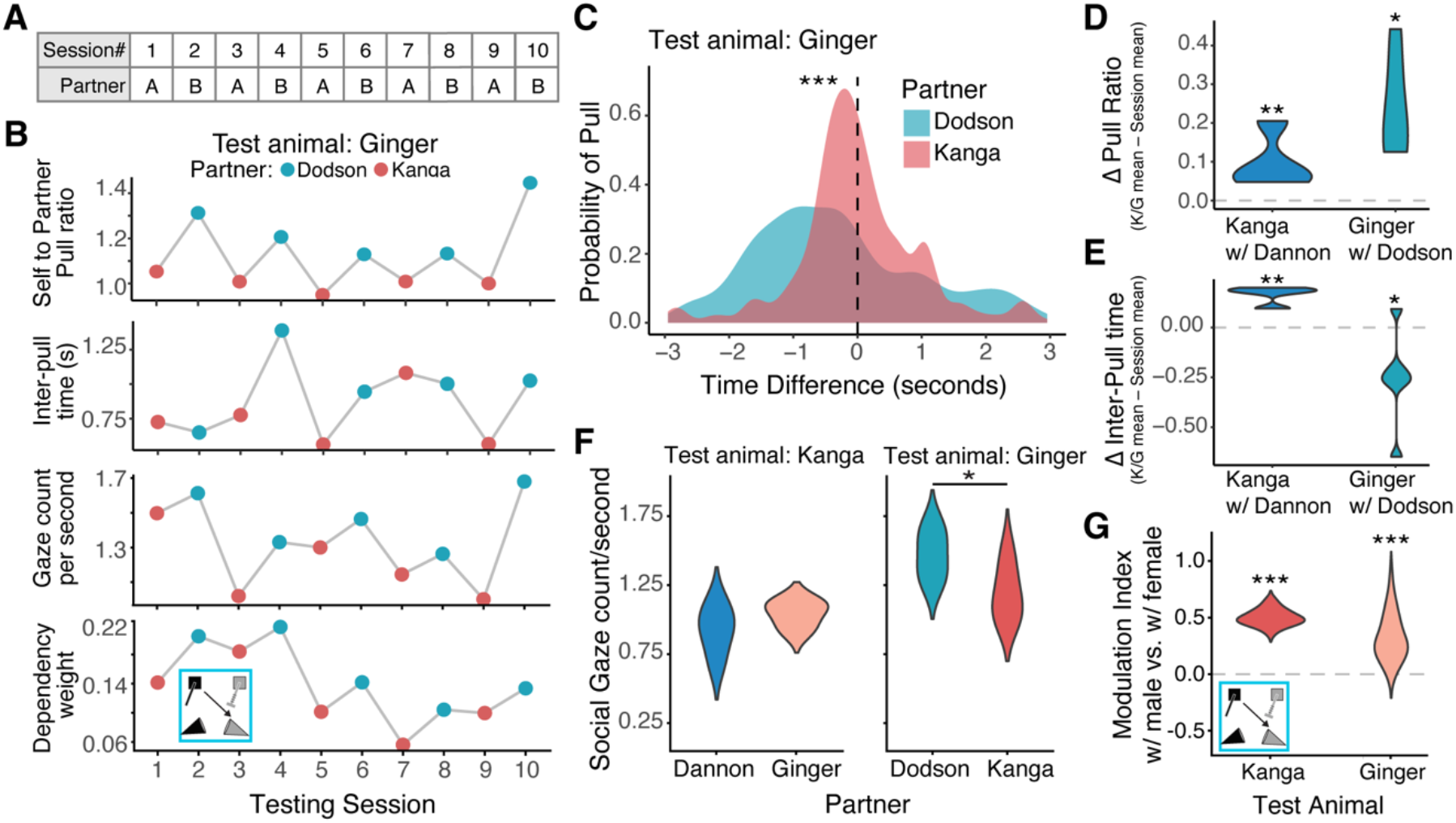
Flexible implementation of partner-specific cooperative strategies. (**A**) Experimental design for testing partner-specific strategies. Two test marmosets completed 10 days of MC sessions, alternating between different partners each day. **(B)** Marmoset’s cooperative behaviors as a function of partner identity across sessions. *Row 1*: Ratio of test marmoset’s lever pulls to partner’s pulls; *Row* 2: mean time interval between test marmoset and partner’s lever pulls; *Row 3*: rate of social gazes directed at partner (gazes per second); *Row 4*: strength of social-attention dependency. (**C**) Probability distribution of test marmosets’ lever pulls relative to their partner’s pull timing, separated by partner identity. (**D**) Comparison of pull ratios between different partner configurations. For each session where test marmosets worked with their pair-mate male partners (Ginger-Dodson, Kanga-Dannon), the mean pull ratio from sessions where Kanga and Ginger worked together was subtracted, showing how pull ratios changed with different partners. (**E**) Comparison of inter-pull times between different partner configurations. For each session where test marmosets worked with their pair-mate male partners (Ginger-Dodson, Kanga-Dannon), the mean inter-pull time from sessions where Kanga and Ginger worked together was subtracted, showing how the timing of coordination changed with different partners. (**F**) Gaze rates for the test marmosets with different partners. (**G**) Modulation index for the test marmosets when cooperating with male versus female partners. Asterisks denote levels of statistical significance (*p < 0.05, **p < 0.01, ***p < 0.001): Mann-Whitney U test with continuity correction for panel C; Wilcoxon signed-rank test for panel D, E and G; Mann-Whitney U test for panel F.

Importantly, partner identity systematically influenced cooperation strategies, with test marmosets showing increased pull ratios when working with their male pair-mate compared to working with each other (Kanga p = 0.065, trend-level; Ginger p = 0.012, t-test; Fig. 6D). Each test marmoset also displayed partner-specific inter-pull intervals (Kanga p = 0.005, Ginger p = 0.045, t-test; Fig. 6E), indicating that they adapted their coordination timing based on their social partner. While Kanga maintained consistent social gaze rates with both partners (p = 0.24, Mann-Whitney U test; Fig. 6F *Left*), Ginger showed partner-dependent modulation of her social gaze (p = 0.049, Mann-Whitney U test; Fig.6F *Right*). Despite this, both Kanga and Ginger showed significantly stronger social-attention dependencies from MTwDBN when cooperating with their pair-mate male partners compared to working with each other, suggesting a greater use of the ‘*gaze-and-pull*’ strategy (p < 0.001 for both animals, Wilcoxon signed-rank test; Fig. 6G). These findings demonstrate marmosets’ ability to flexibly adjust their cooperative behaviors in response to partner identity and social cues. This behavioral flexibility, reflected across multiple measures, underscores the advanced social strategies and cognitive processes employed by common marmosets.

## Discussion

Our study reveals the mechanisms underlying cooperative interactions in common marmosets, demonstrating how these New World primates develop, coordinate, and flexibly deploy social strategies during cooperation. By combining automated behavioral quantification with multivariate causal modeling, we uncovered how marmosets integrate social information and adjust their behaviors to achieve mutual goals, a fundamental aspect of a cooperative society.

The marmosets’ successful mastery of increasingly challenging cooperative timing requirements demonstrated their capacity for behavioral coordination. Importantly, this coordination specifically depended on social vision rather than simple environmental cues or non-visual signals. Consistent with the natural ecology, marmosets dynamically paid attention to each other to monitor what their conspecifics were doing during cooperation (*25, 26*). Their inability to develop alternative strategies in the Automated Lever condition, even after extended exposure, emphasizes that partner-specific social information is fundamental for cooperative success. This reliance on social vision for action planning aligns more closely with human cooperative mechanisms, such as joint attention and gaze-following (*27*), than the olfactory-based social coordination mechanisms seen in rodents (*28*).

During cooperative interactions, marmoset dyads showed sophisticated context-dependent social monitoring strategies. The negative correlation between social gaze and success rates, combined with more precise temporal coupling of gaze during successful versus failed pulls, suggests a nuanced relationship between social monitoring and cooperation. Higher success rates may reflect sessions where dyads achieved better implicit coordination, potentially relying more on peripheral vision or other covert attentional mechanisms rather than direct gaze. When overt social gaze is used, on the other hand, its precise temporal alignment with partner actions during successful pulls indicates that well-timed social monitoring contributes to cooperative success. The increased but less temporally coupled social gaze around failed cooperation suggests marmosets use broader social monitoring as a strategy to re-establish coordination when the behaviors are misaligned with their partner. This strategic deployment of social attention, maintaining precise temporal coordination during successful cooperation while increasing general monitoring to re-synchronize after failures, demonstrates sophisticated social behavioral regulation.

Through dynamic Bayesian network analysis, we examined two distinct cooperative strategies: a ‘*pull-in-rhythm*’ strategy that does not require social gaze and a ‘*gaze-and-pull*’ strategy that integrates social monitoring with action. This pattern of gaze-dependent coordination parallels recent findings in rhesus macaques, where dyads developed transitions from social viewing to action during a cooperative button-pressing task (*14*). By systematically detailing both gaze-dependent and gaze-independent modes of coordination in marmosets, we provide, for the first time, an in-depth characterization of multiple mechanistically distinct cooperative strategies within a nonhuman primate species. Notably, these strategies were specific to the demands of the cooperative task: they did not appear in the self-reward (SR) condition, where pulling was performed independently, nor in the earliest stage of cooperative training (L3), where the 3-second time window was likely too lenient for animals to engage with or learn the true contingency. Their emergence at L2, where the tighter time window required more precise coordination, suggests that these strategies reflect genuine adaptations to the demands of cooperation, rather than artifacts of pulling behavior or general social attention.

The influence of social relationships on cooperation provides crucial insight into how social context shapes cognitive strategy deployment. The emergence of stable leader-follower dynamics, where dominant individuals consistently followed subordinates’ actions, alongside their higher rates of social gaze and lever pulls, suggests that dominant marmosets are more motivated and play an active role in ensuring successful cooperation, consistent with their typically higher food motivation (*29*). This motivation manifests in increased attention to their partner’s actions and more strategic behavioral adjustments to maximize success. Additionally, the superior performance of kin compared to non-kin and mixed-sex compared to same-sex pairs corroborates how pre-existing social bonds and sexual competition shape cooperation (*30*). While kin pairs showed superior performance overall, non-kin mixed-sex pairs demonstrated cooperative ability despite only working together for just three sessions, compared to kin pairs who had months of experience together. This capacity for rapid non-kin cooperation is particularly notable given that cooperation by non-kin is generally considered rare outside of humans (*5*). The apparent scarcity of non-kin cooperation in natural settings may reflect ecological constraints rather than cognitive or motivational limitations (*31*). The alignment of these performance differences with known natural social structures and mating dynamics underscores that performance on our automated cooperative pulling task indeed captures ecologically valid aspects of marmoset behavior.

Most strikingly, marmosets showed remarkable flexibility in adjusting their cooperative strategies based on partner identity, suggesting sophisticated social memory and rapid strategy implementation capabilities. This partner-specific adaptation manifested across multiple behavioral measures, from basic pulling patterns to complex gaze strategies, indicating that marmosets maintain and quickly activate distinct partner-dependent internal models. Their partner-specific cooperative strategies are likely to be shaped by their unique social history, suggesting a sophisticated capacity for social memory, strategy formation, and rapid strategy implementation at the start of each session. This behavioral plasticity, combined with their reliance on social vision and the ability to develop multiple, distinct cooperative strategies, positions marmosets as a valuable model for understanding the evolution of human cooperative cognition (*32*). Altogether, these findings demonstrate that flexible, cognitively guided cooperative strategies extend beyond humans, likely spanning a broader range of the primate order than previously appreciated, with marmosets showing multiple, distinct coordination strategies that they can rapidly tailor based on social relationship factors and history to achieve common goals.

## Acknowledgments

This work was supported by the National Science Foundation Graduate Research Fellowship (DGE2139841, O.C.M.), the National Institute of Mental Health (R21 MH126072, S.W.C.C., A.S.N., M.P.J.), the Simons Foundation Autism Research Initiative (SFARI 875855, S.W.C.C., A.S.N., M.P.J.), Wu Tsai Institute at Yale University (S.W.C.C., A.S.N., M.P.J., W.S.), and a National Eye Institute core grant for vision research (P30 EY026878 to Yale University). We thank Paul Shamble and Joel Greenwood from the Neurotechnology Core of the Kavli Institute for Neuroscience at Yale University and Anthony DeSimone from the Yale School of Medicine Electronic & Machine Shop for providing technical support. We thank Alec Sheffield for consultations on the computational modeling, and Feng Xing on the 3D freely moving animal tracking. We thank the veterinary and husbandry staff at Yale for their excellent animal care.

## Author contributions

Conceptualization: OCM, SWCC, ASN; Experimental Design: OCM, SWCC, ASN, WS; Data Collection: OCM, WS, AN, GN; Data Analysis: WS, OCM, GN; Computational Modeling: WS, MPJ; Supervision: SWCC, ASN, MPJ; Writing: OCM, WS, SWCC, ASN, MPJ

## Methods

### Animals

Six marmosets (three dyads) were first trained on the cooperation task. All pairs were cage-mates: two pair-mate dyads and one mother-daughter dyad. Each marmoset worked exclusively with their partner throughout learning and control sessions. Due to logistical requirements, three trained animals were later re-paired, forming two new pairs: one with a trained and a naive animal, and another with two trained animals who had never worked together as a pair. Data from these new pairs were excluded from the learning analyses (Fig.1) and the analyses of the control conditions (Fig.2), but included in the social gaze (Fig.3) and strategy (Fig.4) quantifications. For testing flexible partner-specific cooperative strategies, four animals (two pair-mate dyads) were used. All marmosets were 2-8 years old, pair- or group-housed, and maintained in the same colony room on a 12-hour light-dark cycle. Before testing sessions, food and water were removed from the cages for 1-3 hours. Upon return to cages after behavior experiments, food and water were given to the marmosets at which point they had unrestricted access. All procedures were approved by the Yale Institutional Animal Care and Use Committee and complied with the National Institutes of Health Guide for the Care and Use of Laboratory Animals.

### Behavioral Experiments: Learning

Marmosets underwent a stepwise training procedure using the MarmoAAP (Marmoset Apparatus for Automated Pulling) (*17*). First, they were trained to enter a transport box and target touch a metal rod for solid rewards (marshmallow or mealworm). After habituation to transport and the testing room, animals were familiarized with transparent behavior boxes and trained to pull levers for liquid reward (marshmallow fluff solution: 6 g fluff/20 ml water).

The initial training consisted of two task condition types: Self-Reward (SR) and Mutual Cooperation (MC). During SR, each marmoset in the pair could independently pull their lever in the presence of one another to receive a 0.1 mL reward, with white squares displayed on the monitor, positioned on the left and right sides, as a visual cue to indicate that the animals are in the SR condition. Once the marmosets consistently performed the SR condition (both animals working continuously for more than 10 minutes to get at least 10 mL of reward per session), they began cooperation training. Unique to the cooperation training phase, we introduced an inter-agent contingency that required both marmosets to pull their levers within a specific time window (cooperation time window, CTW) to earn reward (0.2 mL). To mark the cooperation phase, we displayed two yellow circle cues on the monitor screen to indicate that the animals need to coordinate their pulls. Success was indicated by an immediate auditory tone (2 KHz), followed by reward delivery after a 1s delay. The cooperation time window was initially set at 3.0 seconds (L3) and progressively shortened (3.0s [L3], 2.0s [L2], 1.5s [L1.5], 1.0s [L1]). Dyads advanced to the next phase when they achieved a 50% success rate (proportion of rewarded lever pulls out of all pulls per session) or performed at least 4 sessions. After reaching this criterion in the final 1.0-second window, subsequent sessions were classified as Mutual Cooperation (MC) (Movie S1), representing established performance rather than learning.

### Behavioral Experiments: Controls

After learning the MC condition, dyads underwent a series of control conditions. We first collected two days of baseline MC data. Next, dyads performed No-Vision (NV) control sessions (Movie S2). Marmosets were placed in their transparent chambers and on the apparatus as usual, and then an opaque barrier was inserted between them. This barrier prevented visual contact blocking the views of both the partner and the partner’s lever while leaving auditory and olfactory communication intact. During these NV sessions, marmosets performed the same cooperation condition with the 1-s time window requirement. We ran seven NV sessions to examine whether pairs could adapt to cooperating without visual cues. Following the NV sessions, pairs completed three standard MC sessions to assess whether their cooperative behavior returned to baseline when visual contact was restored.

For the Automated Lever (AL) control condition (Movie S3), we alternated which animal pulled their lever while the other’s lever was fully automated. Sessions were conducted over 12 consecutive days to ensure balanced sampling, with each combination of side (left/right) and action (pulling/automated) repeated three times. The automated lever movements were programmed based on the distribution of lever pull timings from previous MC sessions. The automated lever operated continuously throughout each session, independent of the ongoing behaviors. These sessions maintained the 1-s cooperative time window requirement from the standard condition. Following completion of AL sessions, animals performed at least two standard MC sessions to assess recovery of baseline cooperative performance.

### Behavioral Experiments: Effects of Social Relationships

To evaluate the effect of dominance relationships on cooperative behavior, dominance hierarchies were determined based on established behavioral criteria in marmosets through observations of dyad interactions in their home cage. In marmoset social groups, females typically dominate males, and older individuals dominate younger ones (*33, 34*). Dominant status was confirmed through observation of stereotyped behaviors: dominant marmosets displace others at feeding sites and display characteristic dominant vocalizations, postures, and behaviors, while subordinates respond with deferential behaviors such as yielding feeding priority, moving away from preferred perching spots, and showing ritualized submission postures.

After completing the main cooperation condition and controls, we investigated how social relationships influenced cooperative performance by testing marmosets in novel pairings (number of dyads = 14; F-F = 5, F-M = 6, M-M = 3). We compared the performance of these dyads to that of kin dyads who had previously learned the MC condition together (the original three dyads used for learning analyses). Additionally, we included data from one more kin pair that learned the MC condition after the initial learning experiment for a total of four kin pairs (1 F-F, 3 M-F). We defined non-kin pairs as non-cagemates without direct kinship, though all animals were housed in the same colony room and thus had some degree of familiarity. Each novel dyad underwent a habituation session one day before testing, during which they performed the SR condition. We then aimed to collect three test sessions per novel dyad, though sessions were terminated early or limited to two sessions if animals displayed excessive distress, aggression, or fear behaviors. Standard sessions lasted 20 minutes, and each day multiple pairs of non-kin dyads were tested.

### Behavioral Experiments: Flexible Partner-Specific Social Strategy

We designed a new set of experiments to directly test whether marmosets could flexibly adapt their cooperation strategies based on their partner and their shared history. The study involved four marmosets: two established male-female pair bonds, where the female of one pair was the daughter of the female in the other pair. This daughter provided a valuable test case, as she had first learned the cooperation task with her mother before leaving her family group to form a pair bond with her male partner, with whom she then also learned the task. This arrangement allowed us to study three distinct cooperative relationships: the two pair-bonded dyads and the mother-daughter dyad, where kinship and previous task experience allowed them to easily perform the task despite them no longer being cage mates. We collected data over 10 consecutive days, with the mother and daughter alternating daily between working with their pair-bonded partners and working together, all using the standard MC condition with a 1-s cooperative time window.

### Behavioral Analyses

#### Success Rate

Success rates were calculated for each session by dividing the number of successful lever pulls by the total number of lever pulls for each animal. For AL trials, we excluded pulls made by the automated mechanism and only analyzed the behavior of the pulling monkey. Individual success rates were averaged across both animals to obtain a daily mean success rate per dyad. Success rates of 0 were assigned when no lever pulls occurred. Statistical comparisons between conditions were performed using one-way ANOVA followed by Tukey’s HSD post-hoc tests.

#### Inter-pull Interval

Inter-pull times were calculated by measuring the time interval (in seconds) between sequential lever pulls from each monkey within a trial. To focus on pulls that likely represented intentional cooperative attempts, we only included intervals less than 10 seconds apart (a conservative threshold for identifying attempted coordinated lever pulls). For each session, we calculated the mean inter-pull time across all coordinated pull attempts. These session averages were then compared across learning phases and across experimental conditions (MC, NV, AL) using one-way ANOVA followed by Tukey’s HSD post-hoc tests.

#### Reward Count

For each session, we calculated the total number of rewards earned by each dyad. Daily reward counts were averaged across dyads for each experimental condition. Differences between learning phases and between experimental conditions (MC, NV, AL) were assessed using one-way ANOVA followed by Tukey’s HSD post-hoc tests.

#### Cross Correlation Analyses

To analyze the temporal relationship between animals’ lever pulling behaviors, we performed cross-correlation analyses. Lever pulls were binned into 250-ms time windows across each session, with separate time series created for each animal. For each time bin, we counted the number of lever pulls by each animal. Cross-correlation coefficients were computed for each session always using dominant animals as the reference, with positive lags indicating that subordinate animals’ pulls followed dominant animals’ pulls, and negative lags indicating that dominant animals’ pulls followed subordinate animals’ pulls. We analyzed lags up to ±3 seconds. To assess the significance of temporal correlations, we compared the observed cross-correlation coefficients to shuffled controls where the temporal structure of lever pulls was randomized while maintaining the total number of pulls for each animal. This cross-correlation analysis was performed to assess effects of the control conditions on behavioral coordination performance (Fig. 5A-B) and changes in dyads’ lever pulling coordination across learning (Fig. 2F-G).

#### Event epochs

To explore the temporal relationship among behavioral events, we applied a 3-sec Gaussian kernel to smooth the binary behavioral event variables, defining event epochs (Fig. 3E, *left*). We then correlate the event periods using Spearman correlation (Fig. 3E).

### Markerless Facial Feature Tracking

We used the multi-animal version of DeepLabCut 2.0 (*20*) to track marmoset dyads from the videos taken from three individual cameras (GoPro Hero10) (Fig. 3A). We labeled the animals’ six facial key points, including two ears, two eyes, central blaze, and mouth, that generate the head frame, from 500 frames from five cameras, and trained the DeepLabCut neural network for 150,000 iterations until the error decreased and reached the plateau. Since the three cameras placed in the front of the apparatus were enough to capture most of the animals’ postures and gestures, we only used these three cameras to triangulate the 3D space by using Anipose (*21*). We then defined head gaze as a 15-degree gaze cone perpendicular to the facial plane, estimating the width of marmosets’ viewing angle (*19*), and defined social gaze when one animal’s gaze cone intersects with another animal.

### Vocalization recording and labeling

Vocalizations were recorded using two microphones (Qzt Electronic) placed in the back corners of each transparent box. The audio files were manually labeled using Audacity software. The two audio files from a session were first aligned to session start time, and calls were assigned to specific marmosets by comparing their relative amplitudes across the two microphone tracks - calls with higher amplitude on microphone 1 were assigned to marmoset 1, and vice versa. For each vocalization, the call type was identified according to established marmoset call categories (*35*), and start and end times were marked. For sessions without microphone recordings, audio was extracted from the video cameras and labeled using the same process as above.

To quantify Phee calls during No Vision active and resting periods, we defined rest periods as stretches of time with no lever pulls for at least 10 consecutive seconds. That is, any period with >10 seconds between pulls was classified as a rest period; all other time points were considered active. We then aligned these rest and active time points with the vocalization data to determine whether any Phee calls occurred during each type of period.

### Multi-timelag weighted dynamic Bayesian network (MTwDBN)

To quantify various behavioral strategies, we used an analytic framework that can discover causal dependencies among behavioral variables with minimal supervision across multiple timescales. Dynamic Bayesian Network (DBN) is one such tool for examining the directional dependencies among discrete variables. In particular, a multi-timelag weighted version of DBN (MTwDBN) (*22*) can reveal Granger causal dependencies among social gaze and cooperative pulling behaviors of marmoset dyads. We modified the analysis pipeline as in the previous literature (*22*). We used 1-sec time bins to generate four behavioral variables – the social gaze and lever pull of each animal – and considered three time lags that can discover Granger causalities up to 3 seconds. We applied this MTwDBN pipeline with 3-sec, 2-sec, and 1-sec time lags. To fit MTwDBNs for each task condition (SR, L3, L2, L1.5, L1, MC), we pooled binned behavior data across all sessions of that condition (Fig. 4B,D, S3, S5). To fit MTwDBN for either successful or failed pull conditions during MC, we fit the binned data by ignoring failed or successful pull events, respectively (Fig. 4E). To identify the strategy dependency structure, we used a sampling and bootstrapping technique similar to previous studies (*22*). We performed 200 bootstrap iterations, and in each iteration, we randomly sampled 1,800 data points—matching the smallest sample size across conditions—to avoid bias from sample size differences. We employed the pgmpy Python package (https://github.com/pgmpy/pgmpy) along with custom-written code to determine the best-fitting binary directed acyclic graph (DAG) that captured causal relationships among behavioral variables (real binary DAG). Additionally, we generated a null distribution (null binary DAG) by shuffling time series data and refitting the DBN models. To compute weighted DAGs, we randomly sampled 95 bootstrap iterations, averaged the real and null binary DAGs, and repeated this process 100 times. Across these iterations, we compared real vs. null weighted DAGs to identify significant edges.

To assess how dependencies changed across conditions, we computed modulation indices (∆MTwDBN) by comparing the weighted DAGs of a target condition against a reference condition and normalizing by their sum. In Fig. 4B and D, cooperation conditions (L3, L2, L1.5, L1, MC) were compared to SR. This analysis revealed stable behavioral patterns involving social gaze and pulling behaviors unique to each marmoset during cooperative interactions. In Fig. 4E, we compared DAGs from successful vs. failed pulls in MC to explore strategies linked to successful cooperation. For each marmoset, we averaged ∆MTwDBN values across 100 iterations and pooled results across individuals.

In addition to fitting DBN models, we employed generative DBN models to predict pull actions in MC and assess the significance of different DAG dependency structures (Fig. 4C, 5E). Using the pgmpy package and custom Python scripts, we ran DBN prediction models with 1-sec binned behavioral time series as input, considering a single time lag. We tested the predictive power of four dependency structures: full model – learned structure based on modulation indices between MC and SR; ‘*pull-in-rhythm*’ strategy dependency structure; ‘*gaze-and-pull*’ strategy dependency structure; and combination of both strategies. For each model, we ran 100 iterations, randomly selecting 80% of the data for training. In each iteration, we optimized the generative DBN parameters for the given dependency structure, then tested the trained model on the held-out 20% dataset to predict pull actions. Prediction performance was quantified using Area Under the Receiver Operating Characteristic curve (AUROC/AUC) by comparing predicted and actual pull actions. To account for session-wise variability, we ran this DBN prediction pipeline separately for each session, averaging AUC scores across 100 iterations per session. Final results were reported by pooling across sessions.

## Supplemental Figures and Legends

**Figure S1.**
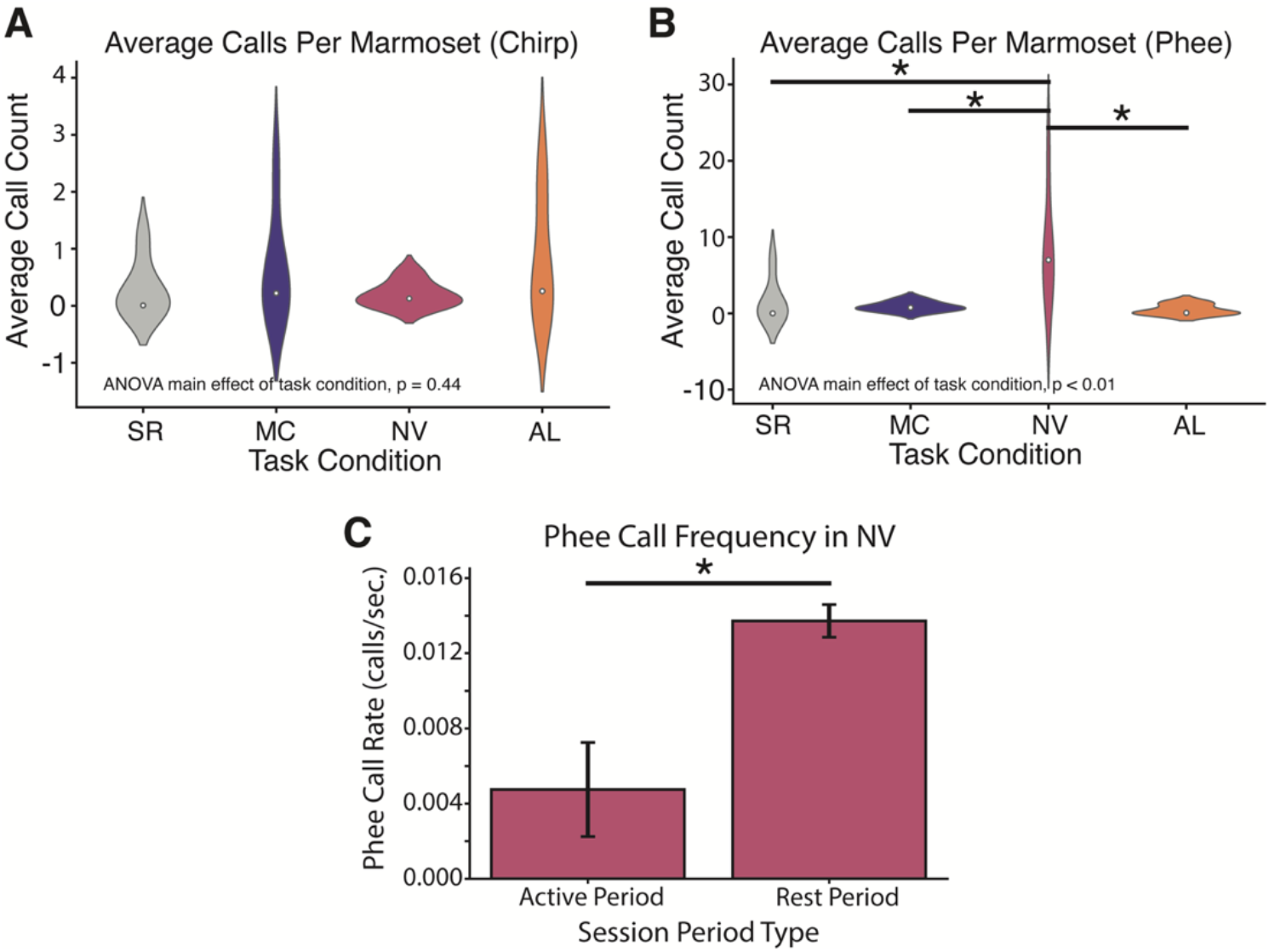
Number of vocalizations in MC and the control conditions (n = 6 animals from 3 dyads). (A) Number of chirp calls during different task conditions. (B) Number of phee calls during different task conditions. (C) Mean phee call rates (calls per second) during active periods (when at least one marmoset pulled the lever within the preceding or following 10 seconds) and rest periods (when no marmoset pulled a lever for at least 10 seconds) in the No Vision task. Asterisks denote levels of statistical significance (*p < 0.05): ANOVA and post-hoc Tukey’s HSD test for panel A and B; Paired t-test for panel C.

**Figure S2.**
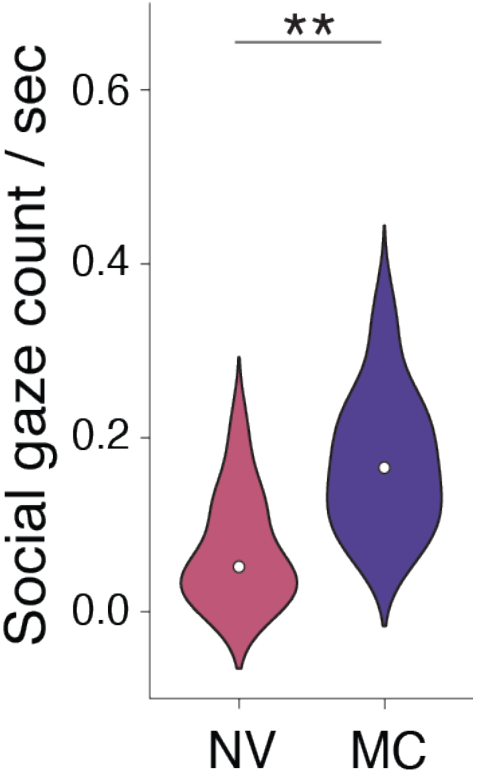
Social gaze count per second in NV and MC (n = 10 animals from 5 dyads). Asterisks denote levels of statistical significance (**p < 0.01): Mann-Whitney U test.

**Figure S3.**
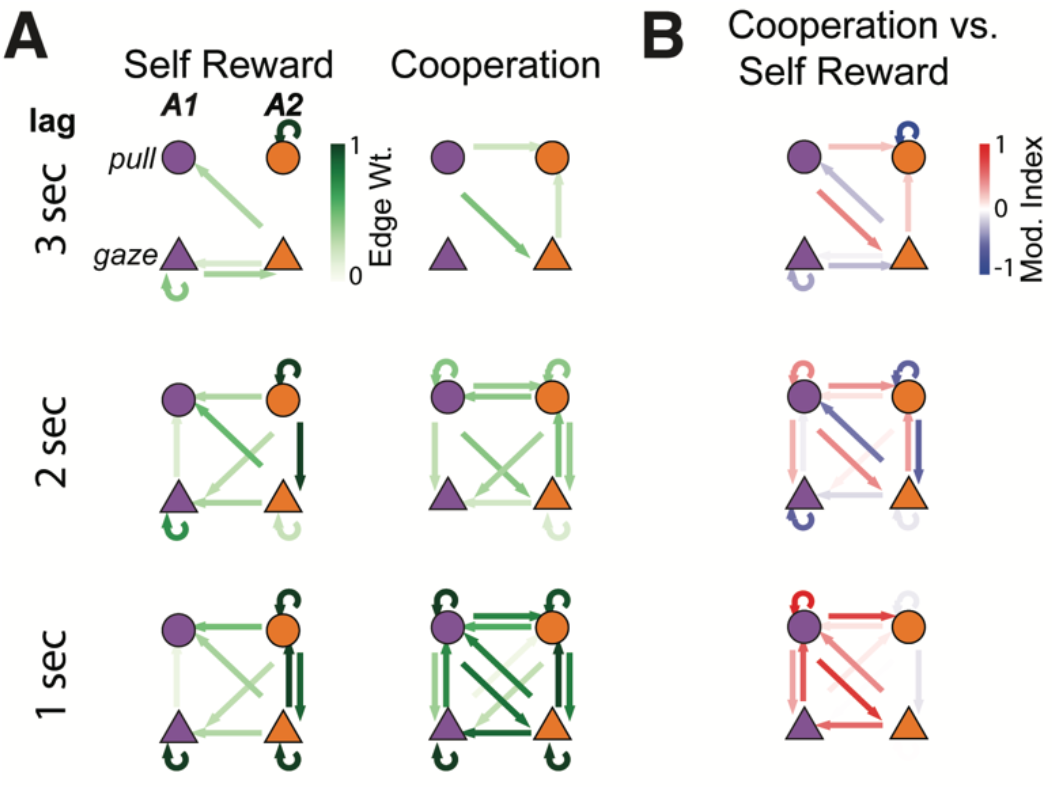
DBN result, example animal pair. (A) Dependencies revealed by three-timelag weighted DBN for three task conditions. (B) the modulation indexes (ΔMTwDBN) between conditions.

**Figure S4.**
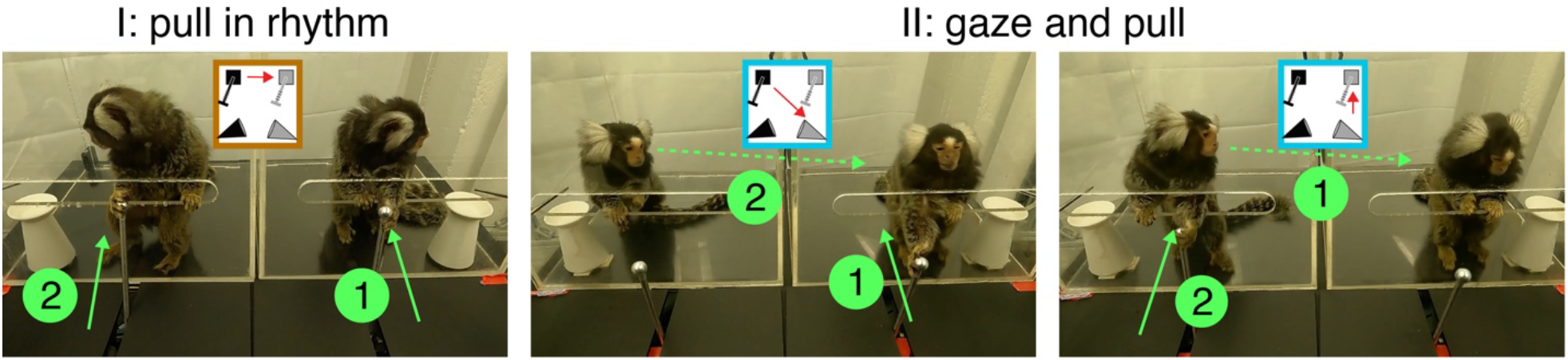
Example frames for *‘pull-in-rhythm’* and *‘gaze-and-pull’* strategies. *Left:* pull-to-pull dependency from the ‘*pull-in-rhythm*’ strategy. *Middle*: social-attention dependency from the ‘*gaze-and-pull*’ strategy. *Right*: gaze-lead-pull dependency from the ‘*gaze-and-pull*’ strategy.

**Figure S5.**
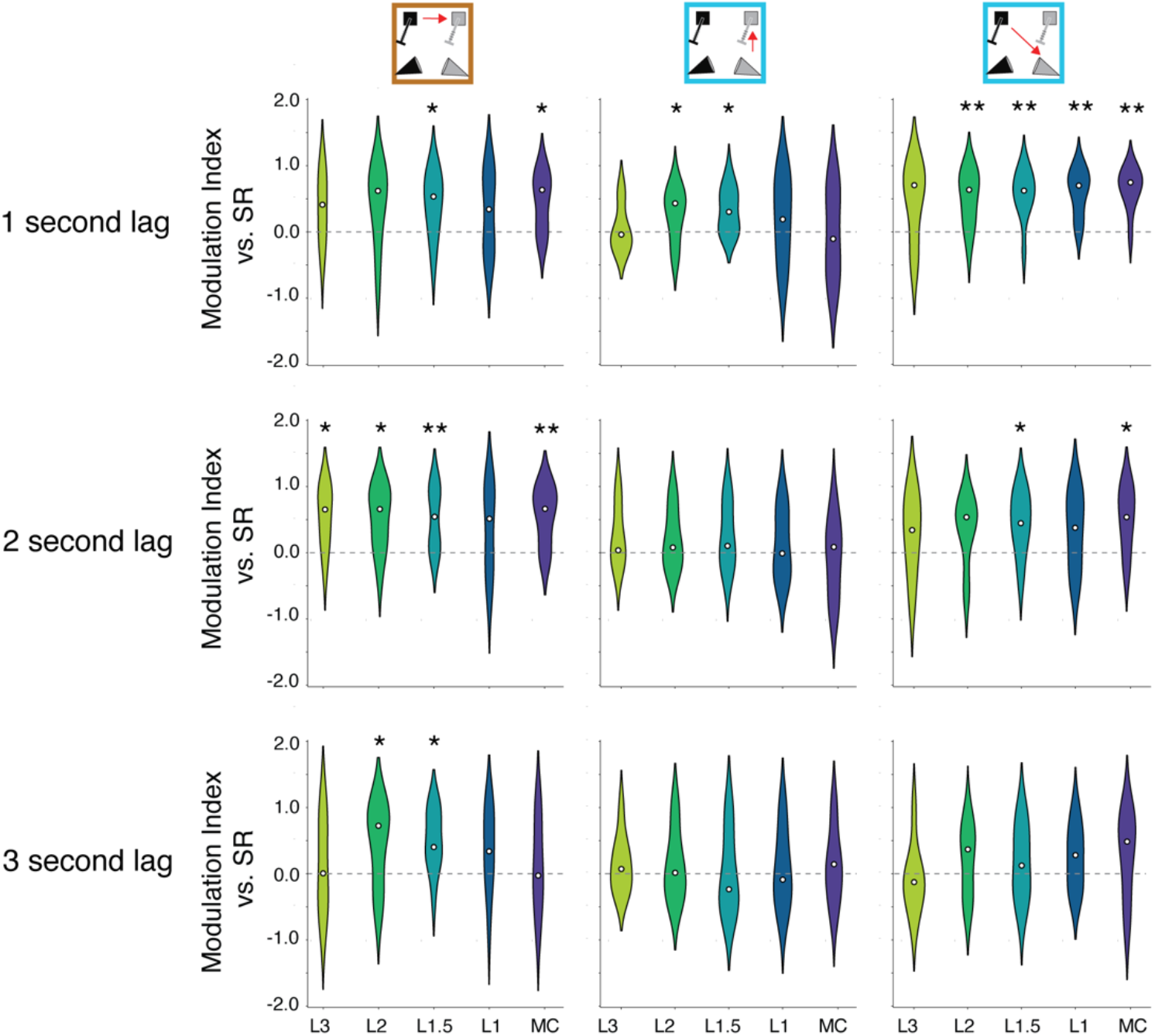
Modulation indices for cooperation conditions with different thresholds across animal individuals, shown three time lags separately. (n = 6 animals for L3 to L1, n = 10 animals for MC). Statistical Significance: Asterisks denote levels of statistical significance (*p < 0.05, **p < 0.01): Wilcoxon signed-rank test.

**Figure S6.**
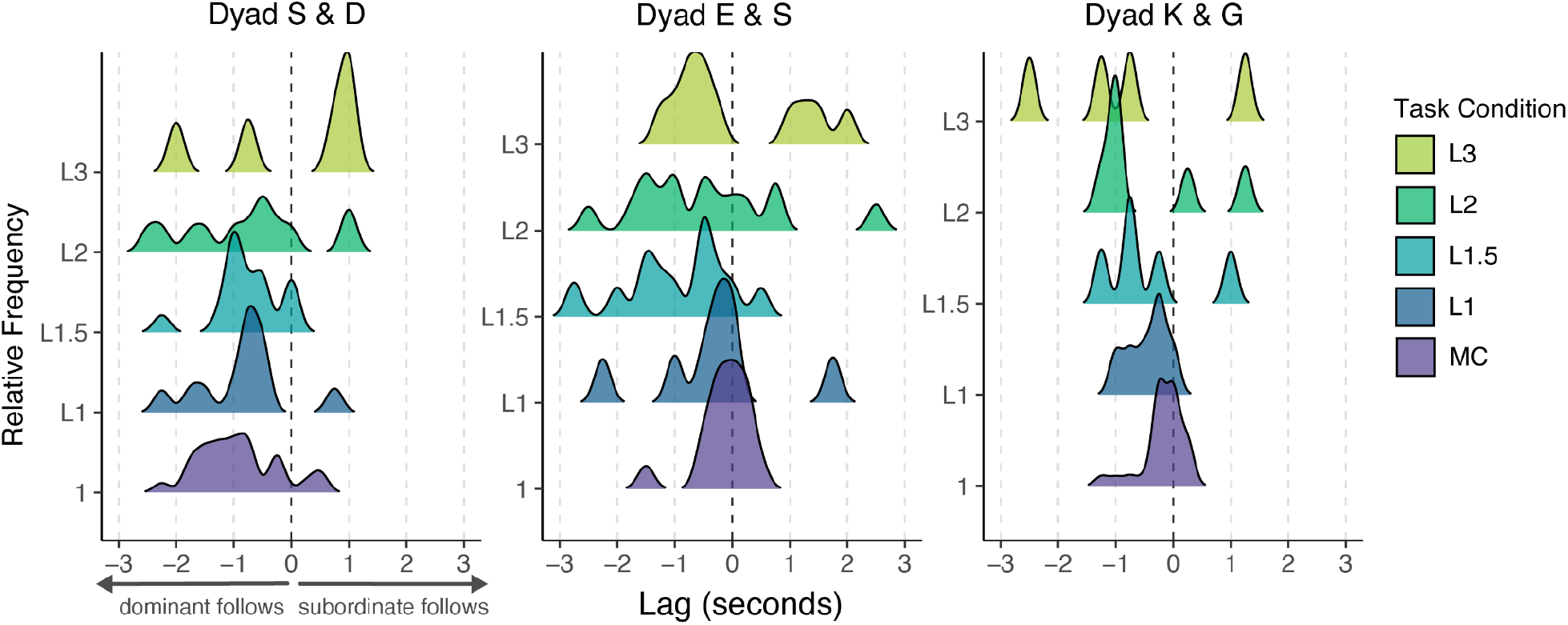
Peak cross-correlation lags across learning for each dyad. Distribution of lag values at peak cross-correlation for each session across training phases. Each subplot represents data from one of the three trained dyads.

**Figure S7.**
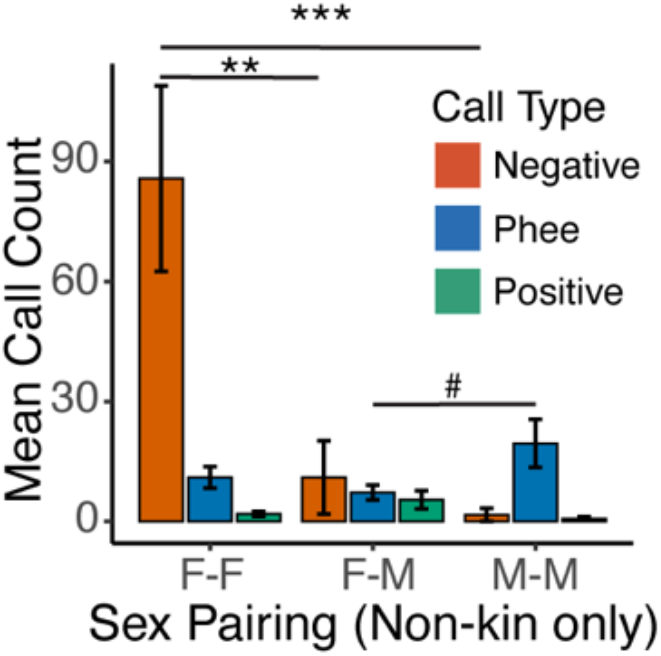
Vocalization of non-kin mixed sex dyads. Non-kin female-female pairs showed more negative calls (e.g., chatter, tsik, ek, tsik-ek), and non-kin male-male pairs showed more phee calls. Positive calls (e.g., trill, chirp) showed no difference among the non-kin pairs. Statistical Significance: Asterisks denote levels of statistical significance (#p < 0.1; **p < 0.01, ***p < 0.001): ANOVA and post-hoc Tukey’s HSD test.

## List of Supplemental Movies

**Movie S1**. Example video clip of 1-second Mutual Cooperation (MC).

**Movie S2**. Example video clip of No-Vision control condition (NV).

**Movie S3**. Example video clip of Automated-Lever control condition (AL).

**Movie S4**. Tracking facial features and defining social gaze.

**Movie S5**. Example clips for the ‘*pull-in-rhythm*’ and ‘*gaze-and-pull*’ strategies.

**Movie S6**. Example video clips demonstrating partner-specific strategies. This movie is played at 1.5 times the original speed.

